# Screening clinical *Candida albicans* isolates for invasiveness by mimicking the human environment

**DOI:** 10.1101/2024.05.27.596042

**Authors:** Clément Vulin, Julian Sutter, Tiziano A. Schweizer, Federica Andreoni, Julian Baer, Alexandra Bernasconi, Karl Bulut, Brunella Posteraro, Maurizio Saunguinetti, Annelies S. Zinkernagel

## Abstract

**Objectives:** *Candida albicans* colonizes a wide range of human body niches, but also causes severe invasive infections, such as candidemia. No screening method exists to predict which colonizer may lead to invasive infections. Invasiveness and virulence are associated with yeast filamentation, triggered by environmental factors encountered in the host. Here, we monitored the filamentation profile and colony appearance time of a *C.albicans* strain isolated from a patient’s abscess. Using eight additional *C.albicans* clinical isolates, we established an *in vitro* screening-framework of filamentation to assess the invasiveness potential of individual isolates.

**Methods:** We monitored the filamentation pattern of nine *C.albicans* clinical isolates over 14 days in 48 environmental conditions, including combinations of glucose/nitrogen concentrations, pH and temperature, to mimic host environment variations. Additionally, we tested invasiveness by growing isolates on modified filtration membranes, mimicking physical human body barriers usually colonized by *Candida*. An automated image analysis pipeline was developed to quantify filamentation.

**Results:** Two types of colony filamentation morphology were differentiated, star and hazy. The total filamentation area depended on environmental factors. Based on their filamentation response to environmental changes, the isolates clustered in three distinct groups reflecting their site of isolation in the host. We moreover found that filamentation morphologies on modified filtration membranes could be predictors of invasiveness.

**Conclusion:** This work lays the ground for screening assays, which could help assessing the potential of a colonizing *Candida* isolate to cause invasive disease, paving the way for tailored preventive therapy regimens in the future.

## Introduction

The yeast *Candida* is a human commensal that can cause invasive candidiasis, mostly in immunocompromised patients[1]. Half of all candidemia cases reported occur in the intensive care unit[2], with *Candida albicans* representing over 50% of the isolated species[3]. Up to 80% of severely ill patients are colonized by *Candida* and the challenge is to predict who will develop invasive disease and therefore profit from early antifungal treatment [4]. Early antifungal treatment can avert severe outcomes in individuals but also drives resistance on a community level[4-6]. Clinical scores in use to decide whether to start early antifungal treatment rely on levels of different biomarkers, risk factors and colonization sites [7, 8]. However, additional isolate-based information could further support clinicians.

*C. albicans* can switch to filamentous forms, characterized by higher virulence and invasiveness than the yeast form[9, 10]. We assessed the filamentation profile and appearance time of colonies formed by a clinical isolate sampled from a patient’s abscess *ex vivo*. As a proxy for invasiveness, we then compared nine *C. albicans* clinical isolates by assessing their *in vitro* filamentation morphology across different nutritional factors (glucose and nitrogen), pH and temperature, and on patterned filtration membranes mimicking host barriers. A machine-learning pipeline was implemented to automate image analysis and to quantify colony filamentation.

## Materials and methods

### Yeast isolates

We processed the patient sample immediately after debridement of the abscess, as previously described[11]. Prior to plating on Columbia 5% sheep blood agar (BioMérieux), the sample was extensively washed and colony growth followed by timelapse imaging for 72 hours at ten minutes time-intervals. A total of nine colonies were recovered for the abscess isolate. Time-lapse image analysis was performed with ColTapp[12]. We analyzed one laboratory strain and nine *C. albicans* isolated from eight patients ([BASEC] No. 2016-00145, 2017-02225, 2017-01140, 2018-01486) admitted to the University Hospital Zürich between October 2017 and November 2019 (**Tab.S1**, **Fig.S1a**).

### Growth conditions

All isolates were grown in yeast-extract-peptone+2% glucose (YPD, BD) overnight shaking at 37°C. 1µL of an overnight culture was inoculated to form a single colony on YPD-agar with variation on pH, temperature, and glucose/nitrogen concentrations for a total of 48 conditions tested (**Tab.S2**, **Fig.S1b-c**). Given that filamentation was uneven on plates with multiple colonies and density-dependent (**Fig.S2**), we inoculated a single colony per plate.

### Colony image acquisition and analysis

Images were acquired using a Canon EOS 1200D reflex camera 3, 5, 7, 9, 12 and 14 days after inoculation (**FigS1d**). ColTapp[12] was used to select colony centers and each colony was extracted in MATLAB (2021b, MathWorks) for further analysis. Segmentation was performed using Ilastik[13] as described in **FigS1e**. Star filamentation is characterized by heterogeneous, distinct filaments, while hazy filamentation by a homogeneous light-grey halo. A final step was performed in MATLAB to correct for filaments mistakenly discovered in the colony center and to extract the colony metrics (colony center radius and maximum filamentation radius) (**Fig.S1f**). Time to first filamentation was obtained from time-series of single colonies by identifying the first day where the maximum filamentation radius diverged (fixed threshold, 100 µm) from the colony radius center. All the codes are available at: github.com/clementvulin/CandidaInvasionScreen.

### Physical barrier membrane model of invasion

Polycarbonate membranes (Millipore, Isopore, 0.2 µm) were modified as previously described[14] by contact printing with polydimethylsiloxane (PDMS). Overnight cultures were inoculated (1 µl) in a central PDMS-free patch (1 mm radius, separated by a 1 mm gap). Quantification of colony spreading was performed by counting the number of patches colonized by *C. albicans* on day 14.

## Results

### *C. albicans* sampled directly from a patient abscess shows a variety of filamentation phenotypes

In a first step, we monitored the growth of a *C. albicans* isolate obtained directly from a patient abscess (**Fig.1a**) and followed the development of colonies. Each colony was assessed for filamentation capacity. We observed that the filamentation phenotype, hazy and/or star (**Fig.S1e**), was not homogeneous within each colony or among colonies (**Fig.1b, Fig.1c**), suggesting either that some phenotypes clustered at the edge of the colony, or that cells switched phenotype during growth, as described previously[15]. We used a 10-minute time resolution to follow colony growth and start of filamentation (**Fig.1d**). The filaments grew faster than the colony center (**Fig.1e**). The initial colony appearance time was very heterogeneous, spanning over 12 hours (**Fig.1f**). Time to first filamentation (hazy+star) also varied among colonies (**Fig.1f**).

**Figure 1:**
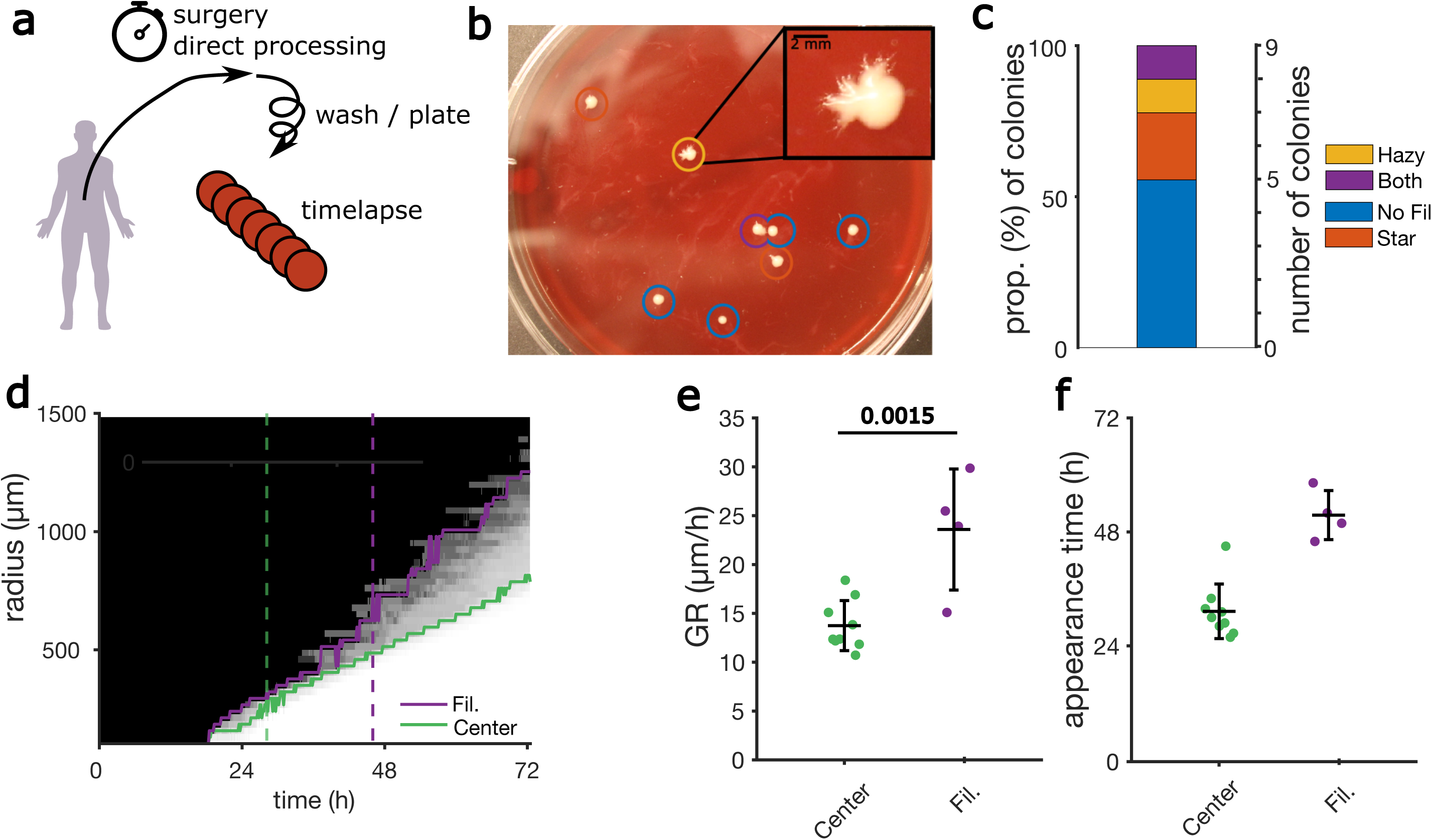
*C. albicans* sampled directly from a patient abscess shows a variety of filamentation phenotypes. **a.** Strain Ab was isolated, immediately after surgery, from a patient presenting with an abscess. Following extensive washing of the debrided material, the yeast was plated on COS and colony growth was monitored over time for 72 hours at time intervals of 10 minutes. **b.** Colonies retrieved after 72 hours of growth. The insert shows a colony displaying asymmetric filamentation. The colored circles denote the filamentation type (yellow=hazy filamentation, red=star filamentation, purple=both types of filamentation, blue=no filamentation. See Fig.S1 and Fig.2 for examples of filamentation types). **c.** Proportion of colonies isolated from the patient sample pertaining to indicated filamentation phenotype after 72h of growth. **d.** Kymograph of the colony depicted in insert **b,** representing the average pixel intensity along the radius and through the time course with center (green) and max filamentation radius (total filamentation, orange) annotated. Vertical dashed lines indicate appearance time (green) and time to first filamentation (orange). **e.** Comparison of the center and filament growth-rates (GR) for each colony. Each dot represents a single colony (only four of them form filament in 72h), **f.** Appearance time of whole colonies (Center) and filament (Fil.), also defined as time to first filamentation. **e.** and **f.** The horizontal bar represents the average and vertical errors bars represent the standard deviation of the mean. The p-value was derived from a two-sample Student’s t-test.

### Filamentation phenotype is isolate dependent

In a second step, we performed an *in vitro* comparative analysis of filamentation profiles of nine *C. albicans* clinical isolates grown in different environmental conditions to replicate the milieu found in the host. Given the right environment, all isolates displayed hazy and/or star filamentation in different proportions, as observed for the patient (**Fig.2a**). However, a multivariable ANOVA with environmental parameters on the colonies’ star filamentation area showed that, starting after 10 days of growth, the strain *per-se* was the parameter explaining most variance (**Fig.2b**).

**Figure 2:**
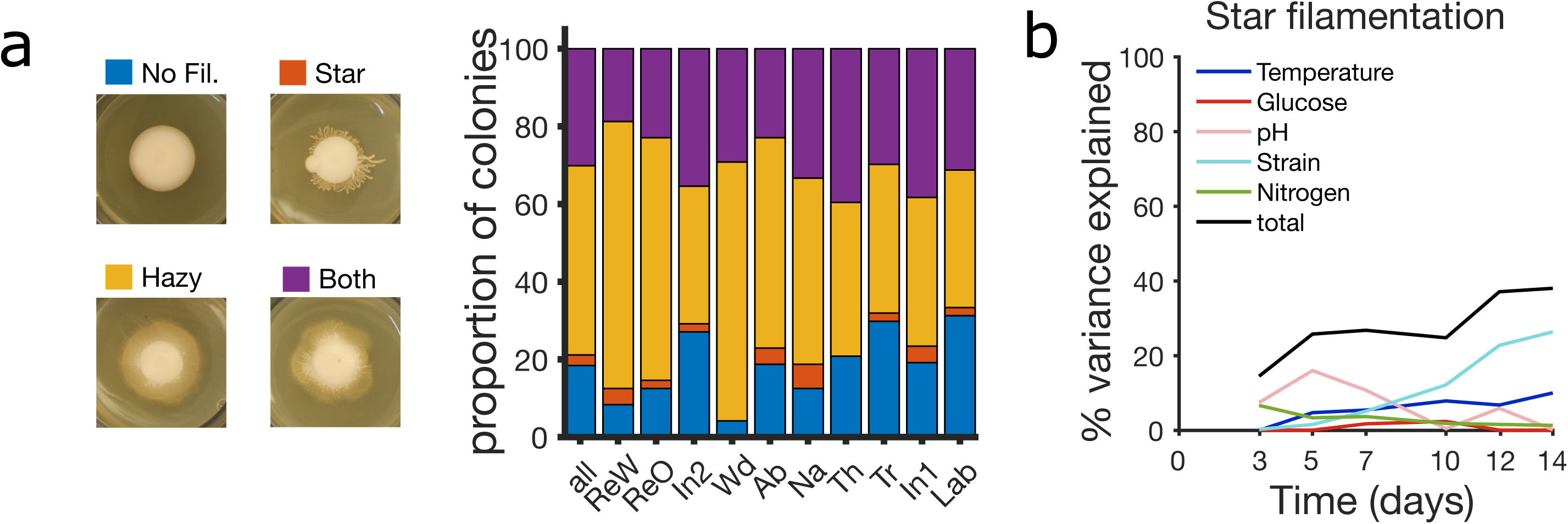
Filamentation phenotype is isolate dependent. Nine *C. albicans* clinical isolates and the Lab reference strain were grown on YPD agar with variations of pH, temperature as well as glucose and nitrogen concentrations, for a total of 48 different environmental conditions tested. Colony growth was monitored over time for 14 consecutive days. **a**. Colonies deriving from the nine clinical isolates displayed heterogeneous filamentation types across all environmental conditions tested (no filamentation (No fil.), star, hazy, or both filamentation types). **b.** Percentage of variance on the “star” filamentation area explained by the different environmental conditions (two biological replicates). Multivariable ANOVA (no interaction terms) for each separate time points across the 48 different environmental conditions (continuous variables: glucose and nitrogen concentration, temperature; categorical variables: pH, strain).

### Filamentation in *C. albicans* is multifactorial and independent from colony growth-rate

In a third step, we assessed the influence of predefined environmental conditions on the total area of filamentation [16]. We found that the filamentation area was dependent on glucose and nitrogen availability for earlier time points (more glucose and less nitrogen leading to more filamentation), explaining cumulatively up to 35% of variance at day 3 (**Fig.3a**). For early time points, initial colony growth rate was not a good predictor of filamentation (**Fig.S3**). In contrast, strain and temperature were the most relevant parameters at day 12 and 14. We therefore choose day 12 as a time point to assess large environmental effects on isolate response and to differentiate isolate-specific filamentation phenotypes and invasiveness potential.

**Figure 3:**
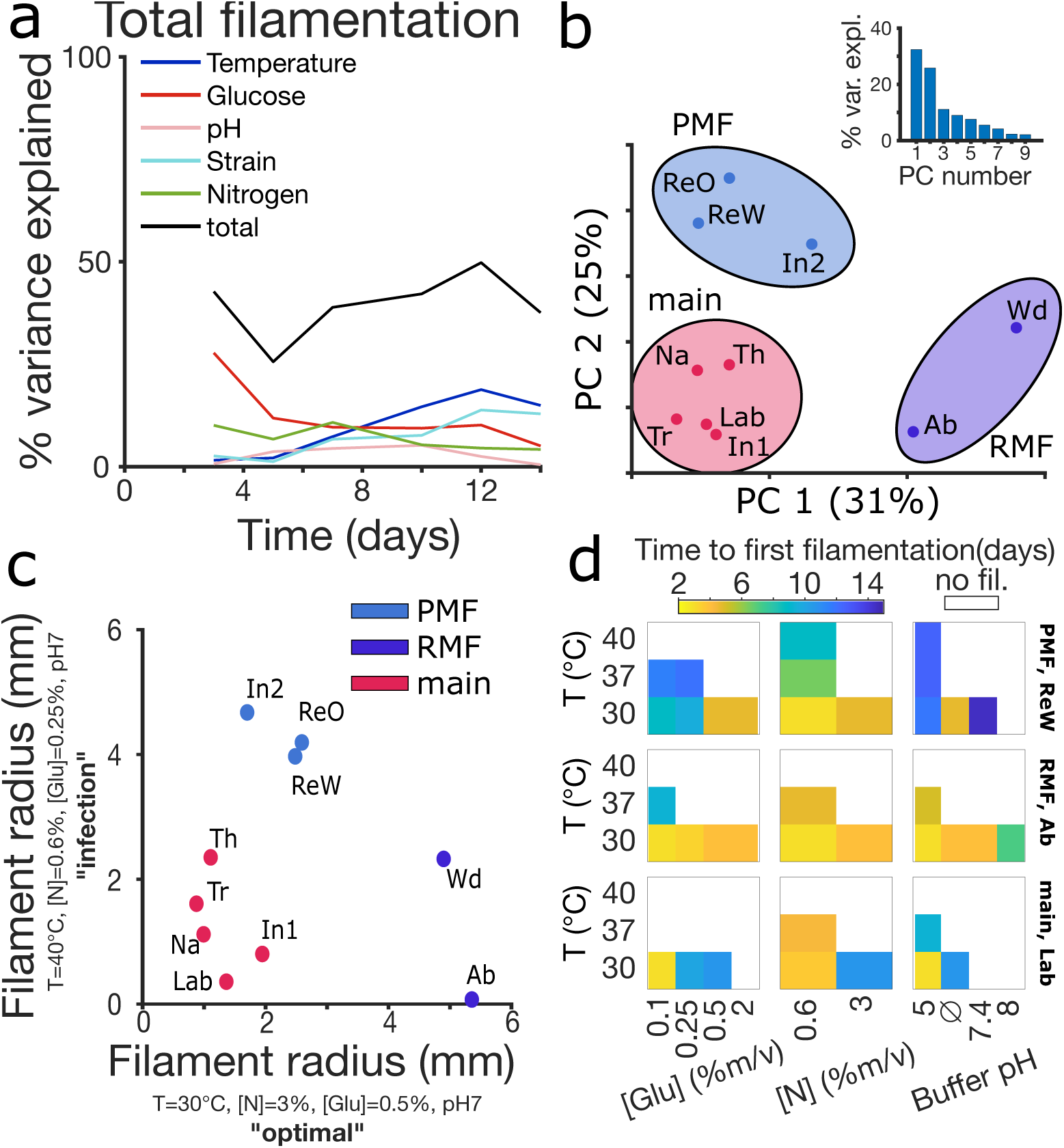
Filamentation is multifactorial. **a.** Percentage of variance on the total filament area explained by the environmental conditions calculated using an ANOVA at different time points of colony growth across 48 different environmental conditions. Colonies were grown for 14 days. Multivariable ANOVAs were performed for each time point separately by considering a linear model without interaction terms with variables pH, glucose and nitrogen concentration, temperature, strains. Explanatory variables were temperature, glucose and nutrient concentrations, pH, and strain. strain and pH were treated as categorical variables. **b.** Principal component analysis (PCA) of the isolates based on their filamentation responses to environmental conditions. The principal component analysis (PCA) considered observation of the average filamentation for each isolate on each of the unique environmental variables. The first two principal components are represented, cumulatively explaining 56% of the variance. The top right insert shows the proportion of variance explained by the principal components (PC). Three groups could be detected, and reflected the isolation site of the strains, invasive (nutrient-poor media filamenters, PMF), superficial (nutrient-rich media filamenters, RMF) and a main group with mucosal isolates (main). **c.** Filamentation response at day 12 using two conditions, “optimal” (Y-axes) or “infection” (X-axes), chosen based on the ANOVA analysis. The grouping found in **b.** could be reproduced. **d.** Average time to first filamentation across conditions. Each row of boxes represents an isolate from one group (ReW for PMF, Ab for RMF, Lab for main), rows within boxes represent temperature. Statistical analysis was carried out using an ANOVA on the environmental parameters and strains (with quantitative and categorical variables, no interaction terms).

We then assessed whether clinical isolates clustered together depending on the amount of filamentation area they produced across the different conditions at day 12 using a principal component analysis (PCA) (**Fig.3b**). Superficial isolates filamented more in nutrient-poor medium (nutrient-poor medium filamenters, PMF) and invasive isolates filamented more in nutrient-rich medium (nutrient-rich medium filamenters, RMF), forming two distinct clusters. The remaining isolates clustered separately and displayed generally lower filamentation (main group). This group included mucosal isolates from the respiratory tract, one superficial isolate and the Lab strain.

Ultimately, to simplify the isolate screening process, we selected two growth conditions that recreated the clustering observed in the PCA (**Fig.3c**), namely “optimal” (30°C, high nutrients) and “infection” (40°C, low nutrients). The main group displayed the lowest filamentation profile, whereas PMFs filamented more under the “infection” condition and less under the “optimal” condition. RMFs filamented more under the “optimal” condition. To assess how quickly filamentation started in a colony, we extracted the time to first filamentation averaged across conditions to assess isolate and group-specific responses (**Fig.3d**). Isolate Re (PMF) displayed long times to first filamentation (7–10 days) while Ab (RMF) displayed shorter times to first filamentation (3–5 days) in all conditions at 30°C. Less filamentation was observed for Lab (main group), especially at high glucose concentrations and alkaline pH. Overall, we observed that filamentation appeared earlier in reduced nutrient concentrations.

### Filamentation allows transmigration of *C. albicans* colonies across physical barriers

We finally tested the isolates for their invasiveness by inoculating colonies on modified filtration membranes[17] presenting a circular pattern of nutrient-rich patches separated by nutrient-devoid gaps, acting as physical barriers (**Fig.4a**) using two conditions (0.1/3 or 2/0.6 % (m/v) of glucose and nitrogen, respectively) favoring either star or hazy filamentation, at two different temperatures (30°C and 37°C). Filamenting colonies managed to transmigrate from patch to patch by stretching filaments across the gaps, denoting their invasiveness (**Fig.4b**). At 37°C, isolates from PMF and RMF had higher invasive capacities than isolates from the main group (**Fig.4c**). We also observed large within-group divergences due to isolate-specific variations.

**Figure 4:**
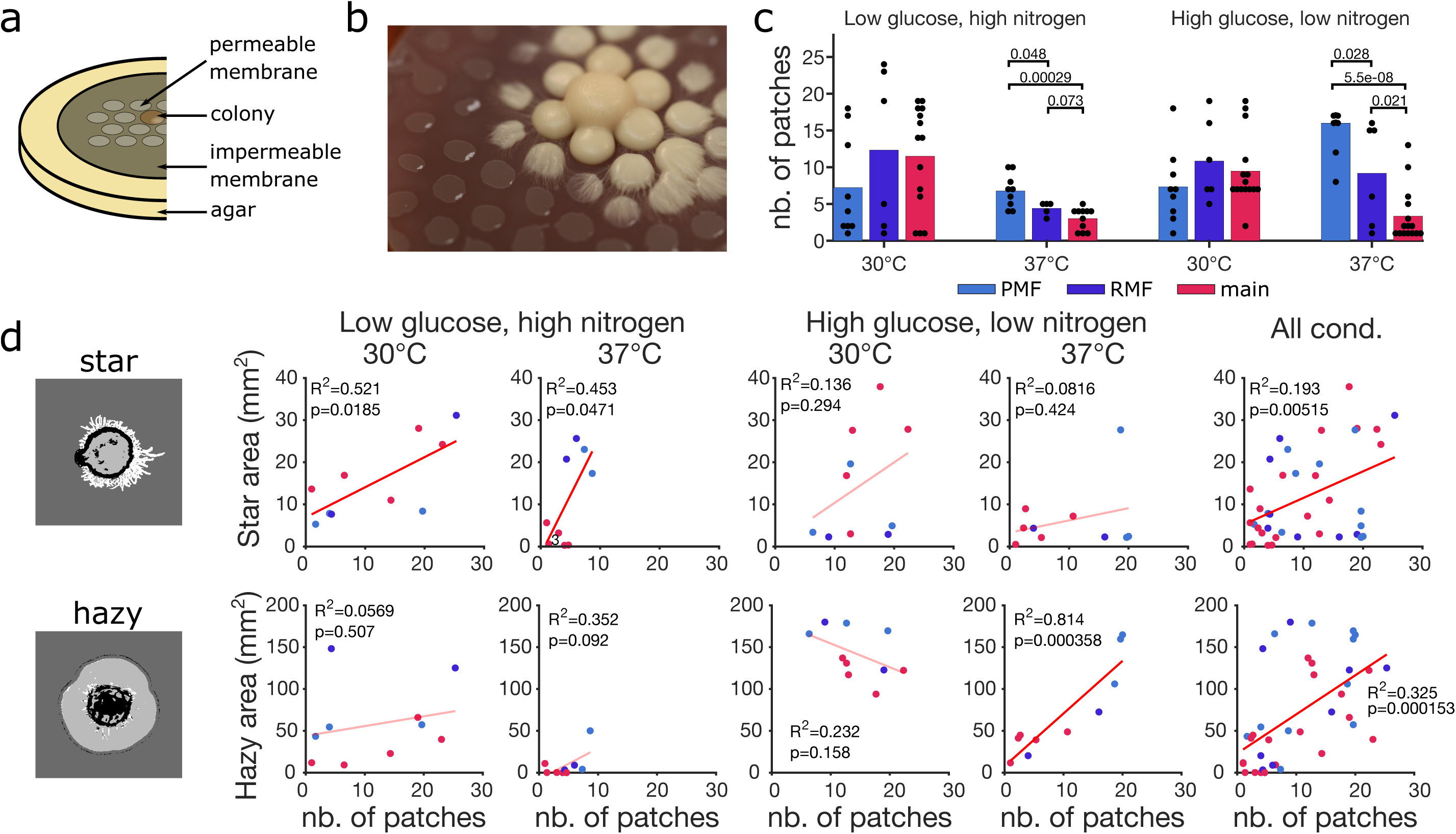
Filamentation allows transmigration of *C. albicans* colonies across physical barriers. **a.** Modified filtration membrane containing circular porous patches (diameter xx mm) that allow nutrients accessibility, separated by nutrient-devoid hydrophobic gaps (PDMS) of distance xx mm. Yeast was initially inoculated in a single, central patch and allowed to grow, spreading across gaps due to the formation of filaments. **b.** Close-up picture of a single colony growing on a membrane, bridging the nutrients-devoid gaps. **c.** Number of invaded patches in four different conditions (low glucose=0.25%, high glucose=0.5%, low nitrogen=0.6%, high nitrogen=3%) after 14 days of growth, for each of the 6 replicates. **d.** Example image segmentation of the star and hazy quantification. Colony center, black; agar, dark grey; hazy filamentation, light grey; star filamentation, white. Comparison between the total number of patches colonized (day 14) and the star or hazy filamentation area (day zz). R^2^ corresponds to the linear regression (red line), p=p-value of the slope parameter. The line is lighter for p-values above 0.05 (not significant).

The amount of star filamentation on agar plates was more predictive for invasiveness in low-glucose/high-nitrogen conditions, while hazy was more predictive in high-glucose/low-nitrogen conditions (**Fig.4d**). This is evident as for the star phenotype, the best fitting linear regressions between the filament area and the number of invaded patches on membranes were observed at low glucose, high nitrogen concentrations regardless of temperature. On the other hand, a significant correlation between the hazy phenotype and invasiveness was only observed in one condition (37°C, high glucose, low nitrogen). In conditions where general filamentation was highest on agar without the membrane (high glucose, low nitrogen, 30°C, **Fig.4d**), some isolates invaded the membrane poorly, which suggested that filament growth with and without nutrients could be de-correlated. Both star and hazy filamentation might explain the ability of a given isolate to spread on the membrane surface when considering all the conditions tested together (**Fig.4d**, “all cond.”).

## Discussion

With this study, we propose a high-throughput screening approach linked to a machine-learning image analysis pipeline to assess the colony filamentation potential of *C. albicans* clinical isolates across environmental conditions and thereby estimate the invasiveness potential of a given isolate, aiming to assist clinicians in the future.

We observed within-colony heterogeneous filamentation phenotypes and a wide range of colony growth-rates, appearance and filamentation times in a *C. albicans* clinical isolate directly sampled from a patient’s abscess indicating that the yeast behavior is dictated by conditions found in the host during invasive disease. This is in line with previous *in vitro* findings[11, 18] describing the phenomenon of persistence. Investigating the causes for the observed filamentation variation could shed light on the behavior of the fungus during infection. In addition to the genetically defined growth patterns, rapid phenotypic variations in yeast are often triggered by hostile environmental conditions encountered in the host [19]. Clinical samples freshly isolated from patients are difficult to obtain and larger cohorts, together with intensive cooperation between physicians and research laboratories, are needed to refine our observations.

We tested variations of pH, temperature and nitrogen/glucose concentrations to model host environment conditions to assess filamentation as a proxy for invasiveness of yeast clinical isolates. An environmental screen of *C. albicans* using six distinct conditions [16] previously showed that filamentation increased in sugar-based media in contrast to amino acid-based or complex media. In this setup, the authors found marginal effects of temperature (30°C and 37°C) on the phenotypes. We used YPD, a complex and nutrient rich growth medium that is not meant to closely represent the human host but allows comparison with a large body of literature. We could show that varying nutrients concentrations and growth conditions using YPD as base medium is enough to trigger filamentation switches, most probably due to nutrient and pH changes occurring during colony growth.

In this work, although colonies grew faster at more alkaline pH, lower temperature, higher glucose or lower nitrogen levels, there were isolate-to-isolate variations and colony growth-rate was not proportional to filament growth-rate. The extent of filamentation was only marginally proportional to the initial growth-rate of the colony. Reflecting what we found in the patient, filamentation was not homogeneous across conditions and heterogeneous within colonies. Two main types of filamentation were observed and in mature colonies the filamentation type depended principally on the isolate tested, indicating a strong genotype effect.

When we considered total filamentation area rather than filamentation type, we found that all conditions taken separately influenced filamentation. Thus, without knowing the exact range of host conditions occurring during an infection, we find it difficult to rank specific conditions as main factors influencing filamentation within-host as suggested previously[16], since the relevance of these variations will depend on the specific host situation. We also expected mixed effects for all parameters tested, and time had a strong influence on the importance of each parameter. Following colony growth for 14 days showed that filamentation, at later time points, depended on temperature (stable throughout the experiment) and on the genotype of the isolate, influencing nutrients absorption rates or production of signal molecules[17, 20]. This suggests that the virulence of a give isolate could change during infection, reflecting changes in filamentation capacity and filamentation type[21-23]. Thus, given the complexity of the potential parameter mapping, future work based on trained machine learning models could be a potential way to predict the invasiveness potential of each isolate. Other less studied factors that typically vary in host conditions, which were not included in this screening set-up, could potentially impact filamentation, such as iron availability[24], interaction with the immune system[25], or the presence of reactive oxygen species[26].

Based on their filamentation patterns, the nine *C. albicans* isolates clustered into three groups that reflected the isolation site. Isolates that originated from superficial sampling (PMF) showed a rather early colony morphology response and a more pronounced filamentation at 30°C and in rich conditions. Invasive isolates (RMF) formed filaments when exposed to infection mimicking environments. Isolates which mainly originated from respiratory tract samples (main group), showed little overall filamentation across conditions. This grouping suggests that filamentation patterns may derive from adaptation to a specific niche characterized by specific environmental conditions, such as high blood glucose levels observed in diabetics, a known risk factor for invasive candidemia[27, 28].

We next adapted this assay to assess the invasiveness of clinical isolates in the laboratory setting by lowering the number of environmental conditions tested from 48 to two, using “infection” or “optimal” conditions-mimicking parameters and quantified filamentation after 12 days of growth, a reasonable time frame for preventive assessments. Using these parameters, we could reproduce the grouping found when all conditions were assessed, opening a new avenue for a diagnostic tool to determine *Candida* invasiveness, implementable in clinical microbiology settings.

The newly developed membrane model also provides a streamlined method for analyzing filamentation patterns, where invasiveness can be assessed rapidly by enumerating the invaded patches. Using specific environmental conditions, we could reproduce the above-described grouping, with the highest filamentation profile associated with invasive isolates. Screening additional isolates will provide further generalizability of this phenomenon. This membrane model could also potentially be used to study the importance of filamentation in the competition with other commensals[29, 30].

Although we observed a general correlation of invasiveness on membranes to filamentation area, we found that not all isolates followed this correlation. Since membranes change the strength of the feedback loop between colony area and how a colony absorbs nutrients based on its diameter, this may be of importance. With membranes, the area of the colony is not necessarily the area in contact with the agar. This means that increasing the size of the initial patch could lead to more favorable conditions for the onset of filamentation. Varying the gap distance between patches to quantify the maximal distance filaments can stretch on a surface without growing from it will further allow measuring the strength of filamentation.

Finally, efforts should be directed towards understanding the changes occurring in the nutritional environment on the human skin as well as within the human body during an infection. Simple tools such as the invasion membrane assays could be used to encourage quantitative measures in laboratories without the need of specific image-analysis expertise. Coupling this assay to larger cohorts tested against media that allow for separation of isolates according to phenotypical behaviors could enhance our understanding of *C. albicans* pathogenicity and lead to better infection prevention strategies.

