## Supplemental material for "Screening clinical *Candida albicans* isolates for invasiveness by mimicking the human environment"

| **Patient ID** | **Strain ID** | **Clinical isolate Nr.** | **Clinical presentation** | **Isolation site** |
| --- | --- | --- | --- | --- |
| 1 | ReW, ReO | 2788  (White, Opaque) | No hospitalisation at the time of sampling | Rectal swab |
| 2 | Th | 3136 | Fournier gangrene | Throat swab |
| 3 | Na | 4156 | Fistula dorsal neck (CSF fistula, prosthesis-associated infection) | Nasal swab |
| 4 | In1 | 4132 | Elective surgery to reconnect the bowel | Inguinal swab |
| 5 | In2 | 3943 | Infected seroma | Inguinal swab |
| 6 | W | 3288 | Sealed rupture of mycotic aneurysma left arteria iliaca externa | Deep-seated wound |
| 7 | Tr | 2439 | Vascular graft infection, pacemaker and prothestic valve endocarditis | Tracheal secretion |
| 8 | Ab | 1269 | Abscess | Abscess pus |
|  | Lab | ATCC 90029 | NA | Blood |

**Supplementary table 1: Patients and strains information**

Eight patients were included in this study. Three strains were isolated from the respiratory tract (nasal swab, “Na”; throat swab, “Th”; tracheal secretion, “Tr”), two strains from inguinal swabs (“In1” and “In2”), one from a rectal swab (“Re”), and two strains were invasive isolates (abscess pus, “Ab”; deep-seated wound, “Wd”). The strain isolated from the rectal swab presented as a white or opaque variations (“ReW” and “ReO”, respectively), and we performed our experiments on cultures started separately from white or opaque colonies. CSF=cerebrospinal fluid, NA=not available

| **Condition** | **Temperature (ºC)** | **Glucose**  **(% w/v)** | **Nitrogen**  **(% w/v)** | **pH** |
| --- | --- | --- | --- | --- |
| 1 | 30 | 0.1 | 0.6 | no buffer (7) |
| 2 | 30 | 0.1 | 3 | 5 |
| 3 | 30 | 0.1 | 3 | 7.4 |
| 4 | 30 | 0.1 | 3 | 8 |
| 5 | 30 | 0.1 | 3 | no buffer (7) |
| 6 | 30 | 0.25 | 0.6 | no buffer (7) |
| 7 | 30 | 0.25 | 3 | 5 |
| 8 | 30 | 0.25 | 3 | 7.4 |
| 9 | 30 | 0.25 | 3 | 8 |
| 10 | 30 | 0.25 | 3 | no buffer (7) |
| 11 | 30 | 0.5 | 0.6 | no buffer (7) |
| 12 | 30 | 0.5 | 3 | 5 |
| 13 | 30 | 0.5 | 3 | 7.4 |
| 14 | 30 | 0.5 | 3 | 8 |
| 15 | 30 | 0.5 | 3 | no buffer (7) |
| 16 | 30 | 2 | 3 | no buffer (7) |
| 17 | 37 | 0.1 | 0.6 | no buffer (7) |
| 18 | 37 | 0.1 | 3 | 5 |
| 19 | 37 | 0.1 | 3 | 7.4 |
| 20 | 37 | 0.1 | 3 | 8 |
| 21 | 37 | 0.1 | 3 | no buffer (7) |
| 22 | 37 | 0.25 | 0.6 | no buffer (7) |
| 23 | 37 | 0.25 | 3 | 5 |
| 24 | 37 | 0.25 | 3 | 7.4 |
| 25 | 37 | 0.25 | 3 | 8 |
| 26 | 37 | 0.25 | 3 | no buffer (7) |
| 27 | 37 | 0.5 | 0.6 | no buffer (7) |
| 28 | 37 | 0.5 | 3 | 5 |
| 29 | 37 | 0.5 | 3 | 7.4 |
| 30 | 37 | 0.5 | 3 | 8 |
| 31 | 37 | 0.5 | 3 | no buffer (7) |
| 32 | 37 | 2 | 3 | no buffer (7) |
| 33 | 40 | 0.1 | 0.6 | no buffer (7) |
| 34 | 40 | 0.1 | 3 | 5 |
| 35 | 40 | 0.1 | 3 | 7.4 |
| 36 | 40 | 0.1 | 3 | 8 |
| 37 | 40 | 0.1 | 3 | no buffer (7) |
| 38 | 40 | 0.25 | 0.6 | no buffer (7) |
| 39 | 40 | 0.25 | 3 | 5 |
| 40 | 40 | 0.25 | 3 | 7.4 |
| 41 | 40 | 0.25 | 3 | 8 |
| 42 | 40 | 0.25 | 3 | no buffer (7) |
| 43 | 40 | 0.5 | 0.6 | no buffer (7) |
| 44 | 40 | 0.5 | 3 | 5 |
| 45 | 40 | 0.5 | 3 | 7.4 |
| 46 | 40 | 0.5 | 3 | 8 |
| 47 | 40 | 0.5 | 3 | no buffer (7) |
| 48 | 40 | 2 | 3 | no buffer (7) |

**Supplementary table 2: List of condition tested in this study**

The table lists the 48 conditions tested to assess filamentation changes in the yeast *Candida albicans*. Temperatures of 30, 37 or 40ºC were testes as well as glucose concentrations of 0.1%, 0.25%, 0.5% and 2% (%w/v) and nitrogen concentrations of 0.3% and 6% (%w/v). YPD agar was also used either unbuffered (starting pH value of 7) or buffered, to obtain a pH of 5.5, 7.4 or 8. YPD agar was buffered to pH5.5 with citrate and Na_2_PO_4_, to pH7.4 with Na_2_H_2_PO_4_ (0.4M) and to pH8 with 1M Tris. For each strain three colonies were assessed, the experiments were repeated twice.

**Figure S1: Strains origin and experimental setup*.***

**a.** Map of strains recovered from different body sites. Two strains were collected from invasive infections (Wd, Ab), three from inguinal and rectal swabs (In1, In2, ReW and ReO), three from the respiratory tract (Na, Th, Tr). The colors correspond to the principal component analysis (PCA) groupings in Fig.3: red: main group, dark blue: nutrient-rich medium filamenters (RMF) group, light blue: nutrient-poor medium filamenter (PMF) group. **b,c.** We assessed 48 different combinations of culture conditions based on YPD medium with temperature, pH and glucose variations at **b.** low (06%) and **c.** high (3%) nitrogen levels (Z-axis in **c.**, “no B”: unbuffered condition). **d.** Each colony was followed over time for 14 days, to determine colony center radius (green circle) and maximum filamentation radius (orange circle). The example pictures refer to the following conditions for strain Ab: 37°C, [Glu]=0.1, %m/v, [N]=1, pH=7. **e.** Examples of analysis for detection of colonies characteristics and features. Represented are three typical phenotypes, no filamentation (no fil.), “hazy” filamentation (homogeneous thin filaments, corresponding to single hyphae spreading close to the surface of the agar) and “star” filamentation (thicker, venous filaments up to 0.5 mm across). After automated discovery of colonies (“ori.”: originals), a two-step machine learning algorithm was used. Segmentation was performed with the Ilastik software trained on 26 representative images of colonies. Each colony was first segmented from the background followed by sorting of the colony center from filaments, classifying pixels into 4 groups: colony center, black; agar, dark grey; hazy filamentation, light grey; star filamentation, white. Finally, segmentation of the colony center was refined using geometrical rules (“final”). **f.** Example of colony and filament growth (37°C, [Glu]=0.1 %m/v, [N]=1, pH=7, strain Ab), showing the measured colony center radius (green) and maximum filamentation radius (orange). Each dot represents one replicate, and the vertical dashed line is the calculated mean time to first filamentation. For each strain three colonies were assessed (technical replicates), the experiments were repeated twice.

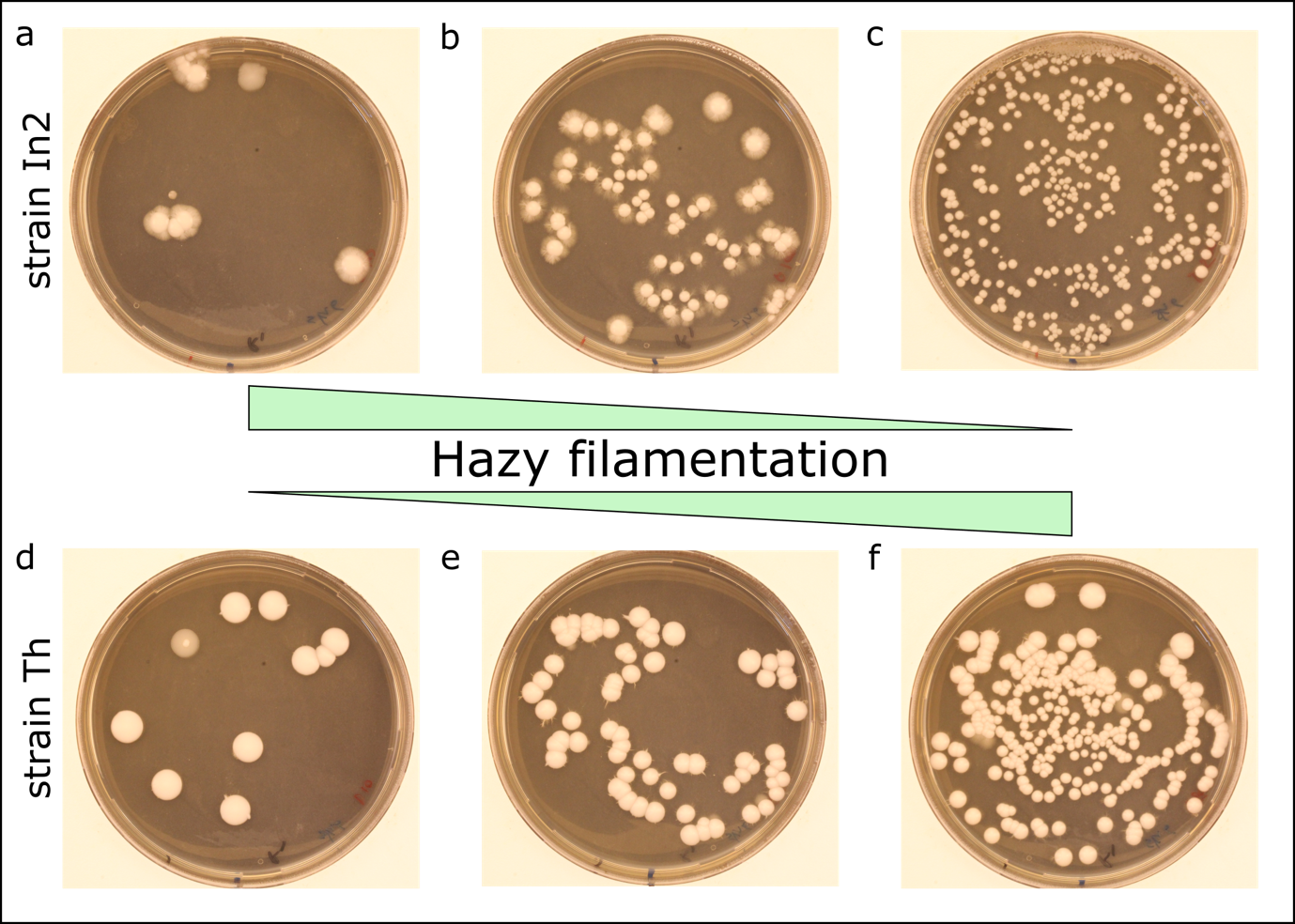

**Figure S2: Effect of density on filamentation and phenotypic variation in colonies derived from single cells.**

When plating strains (panel **a-c** strain In2, panel **d-f** strains Th) at different colony forming unit (CFU) numbers per plate, we observed density dependence of filamentation. Moreover, while strain In2 filaments less at higher density, strain Th filaments differently. The most convincing effect of density can be observed in panel b, where filamentation is only observed for larger colonies at the less dense periphery of the plate. In panel f “Star” filamentation and “Hazy” filamentation occur most prominently side by side. As filamentation was uneven on plates with multiple colonies and density-dependent, as shown above, subsequent experiments were were carried out assessing filamentation of a single colony.

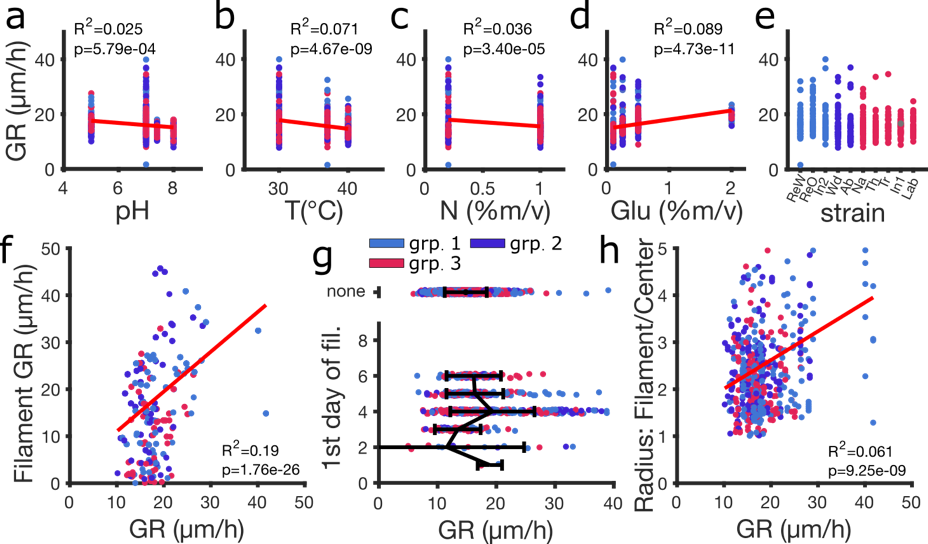

**Figure S3: Filamentation is independent from colony growth-rate.**

**a.-d.** Effect of environmental factors (pH, temperature (T), nitrogen (N) and glucose (Glu) concentration) on colony growth-rate. **e.** Colony growth-rates observed for each strain across all conditions. **f.-h.** Effect of growth-rate on **f.** filament growth-rate, **g.** first day of filamentation and **h.** the proportion of colony radius occupied by filaments at day 14 (ratio of radii). Dot colors correspond to groups as determined in Fig.3b. The R^2^ correspond to a linear regression (red line), the represented p-value (p) is testing against the null-hypothesis that the slope coefficient is different from zero. The black line in **g.** links the means of each group and error bars represent the standard deviation of the mean. Fil.: filamentation. GR: growth-rate.
